# Higher multiplication rates of *Plasmodium falciparum* in isolates from hospital cases compared with community infections

**DOI:** 10.1101/2024.05.02.592253

**Authors:** Lindsay B. Stewart, Elena Lantero Escolar, James Philpott, Antoine Claessens, Alfred Amambua-Ngwa, David J. Conway

## Abstract

**Background:** Parasite multiplication rates vary among *Plasmodium falciparum* isolates from patients with malaria, suggesting differences in virulence potential, although direct comparisons between hospital-based clinical cases and community infections are needed.

**Methods:** Cryopreserved blood samples from malaria cases presenting to a district hospital in The Gambia and infections detected in local communities were introduced to continuous culture under the same conditions. Thirty-four isolates (23 hospital-based and 11 community-based) were successfully established and then tested under exponential growth conditions over six days to derive estimated *P. falciparum* multiplication rates per cycle based on a 48-hour typical cycle length.

**Results:** A range of parasite multiplication rates in culture was seen across isolates, from 1.5-fold to 5.0-fold per cycle. Multiplication rates were significantly higher in the hospital-based isolates than the community-based isolates. There was a significantly positive correlation between parasitaemia in peripheral blood and multiplication rates in culture. There was no significant difference in multiplication rates between isolates with single or multiple parasite genotypes.

**Conclusions:** These findings are consistent with a hypothesis that intrinsic natural variation in parasite multiplication rate may affect levels of parasitaemia achieved during infection, and that this affects likelihood of hospital presentation. Results do not support a hypothesis that parasites modify their multiplication rates in response to competing parasites with different genotypes.

**Summary:** Relevant to understanding parasite virulence, this study finds higher *Plasmodium falciparum* multiplication rates in cultured isolates from malaria cases presenting to hospital than in isolates from local community infections, and positive correlation with parasitaemia in peripheral blood of individuals.

## Introduction

All clinical malaria is caused by asexually replicating blood-stage parasites, with high infection loads leading to severe disease in some cases [1, 2]. Despite previous advances in malaria control, the global burden of disease remains very high and extremely inequitable, more than half of all annual malaria cases and deaths being due to *Plasmodium falciparum* in West Africa [3]. Any variation in parasite multiplication potential would be likely to influence infection load and disease risk, so its importance needs to be determined.

Parasite multiplication rates cannot normally be measured directly in patients, as therapy requires prompt anti-parasite treatment, although modelling of multiple parameters in clinical samples suggests there is intrinsic variation of parasite multiplication potential [2, 4]. Moreover, it is challenging to estimate parasite multiplication rates in a standardised manner in induced experimental infections with laboratory strains of *P. falciparum* [5]. Interestingly, one study suggested a multiplication rate difference between two unrelated *P. falciparum* laboratory strains [6], although another independent study elsewhere did not indicate a difference after a similar comparison [7]. Interpretations from experimental infections are also complicated by the fact that genomic changes which are not seen in natural infections have occurred within laboratory strains [8].

Importantly, studies of parasites in short-term cultured clinical isolates have indicated variation in multiplication rates which would not be affected by mutants that may arise after long-term culture. Measuring *P. falciparum* multiplication in the first cycle *ex vivo*, studies in Thailand and Uganda [9, 10] showed higher rates in severe malaria compared to mild malaria isolates, although a study of parasites from Mali and Kenya indicated no difference between severe and mild malaria isolates [11]. Although interesting, a limitation of these comparisons is that initial viability of parasites within clinical samples is generally variable, affecting interpretations of multiplication in the first *ex vivo* cycle.

Malaria parasite multiplication rate measurements can be more controlled under conditions of continuous *in vitro* culture. After establishment in continuous culture for a few weeks, under exponential growth conditions the multiplication rates of *P. falciparum* in diverse clinical isolates have shown a range from approximately 2-fold to 8-fold (per 48-hour period corresponding to a typical cycle) [12, 13]. Interestingly, among isolates from a highly endemic population in Ghana, there was a significantly positive correlation between exponential multiplication rate in culture and the parasitaemia in peripheral blood of patients [13], suggesting that *in vitro* assay measurement correlates with multiplication potential *in vivo*. Such multiplication rate variation among newly-established clinical isolates is not affected by parasite mutants that may become common in longer-term cultures, which affect the most widely-cultured and compared laboratory-adapted strains of *P. falciparum* [8, 14].

To investigate naturally occurring parasite multiplication rate variation, it is also important not only to study isolates from clinical cases, but also from infections detected within local communities, as some of these might contain parasites with lower intrinsic multiplication rates than those that cause clinical malaria. Although such community infections are common, it is generally harder to study parasites from these as they often have very low levels of parasitaemia [15]. This study presents the first comparison of multiplication rates between parasites from hospital cases and from infections sampled in the local community. Malaria In The Gambia remains endemic, with almost all cases being caused by *P. falciparum*, although the incidence is lower than experienced two decades ago [16-19]. To analyse intrinsic multiplication rates of parasites, isolates from cryopreserved blood samples of hospital cases and community infections have been newly established and assayed in under exponential growth conditions in culture. The potential importance of intrinsic multiplication rate variation is suggested by higher multiplication rates in the hospital isolates, and a positive correlation between parasitaemia in peripheral blood and multiplication rates in culture.

## Results

### P. falciparum multiplication rates in cultured isolates from hospital and community samples

Cryopreserved blood samples from malaria cases presenting at Basse Hospital and local community *P. falciparum* infections were thawed and cultured under the same conditions for at least 20 days prior to testing in six-day exponential multiplication rate assays, yielding high quality data for 34 isolates (Figure 1A, Supplementary Tables S1 and S2). Community-based isolates (N = 11) showed a low range of multiplication rates from 1.5-fold to 2.9-fold (expressed as the fold-change per 48 hours corresponding to a typical cycle length). The hospital-based isolates (N = 23) showed a range from 1.5-fold to 5.0-fold.

**Figure 1.**
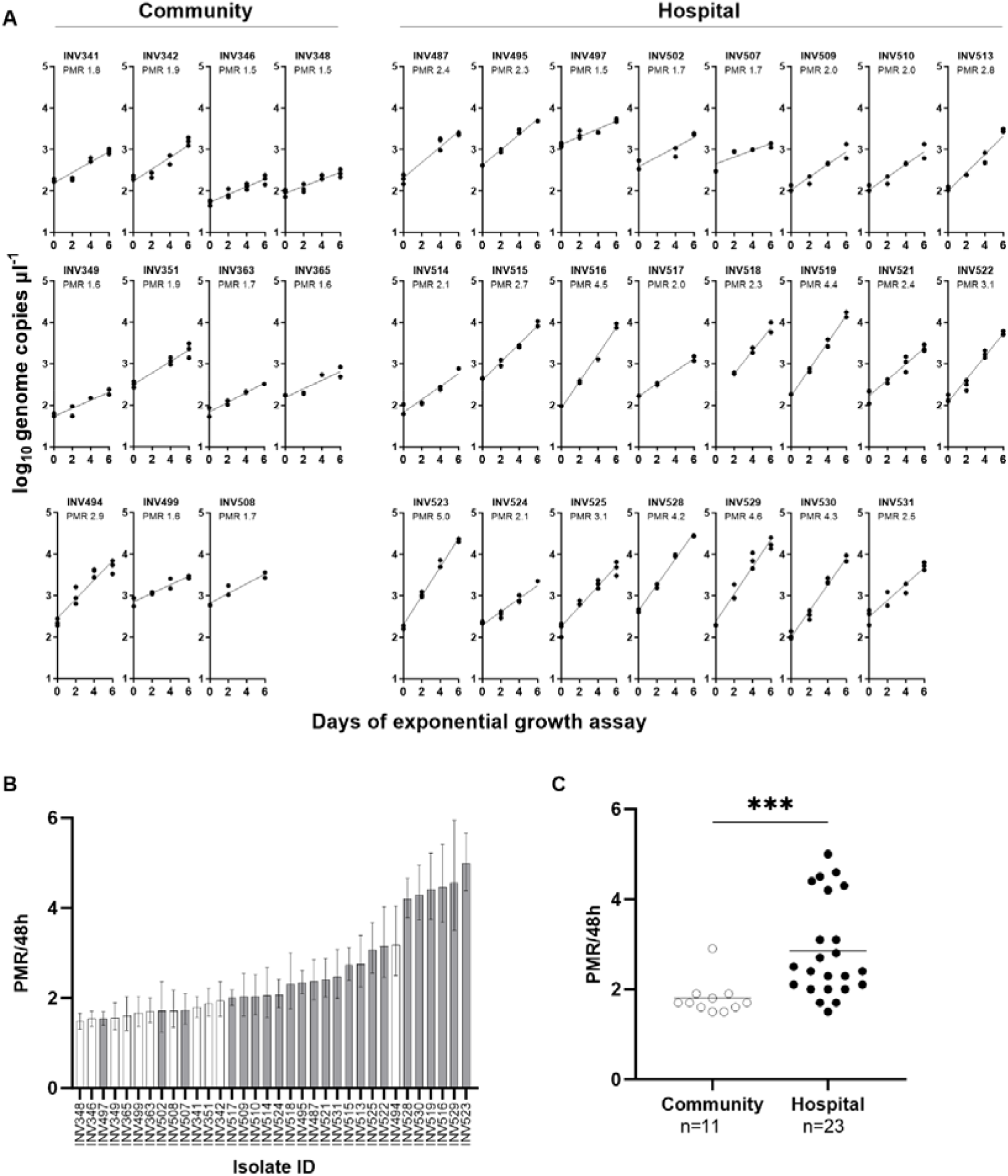
**A**. Exponential multiplication rate assays of 34 Gambian *P. falciparum* isolates. Community-based isolates (N = 11) are shown on the left, and hospital-based isolates (N = 23) on the right. After at least 3 weeks of continuous culture, each isolate was tested in a 6-day exponential growth assay conducted in triplicate with erythrocytes from three different donors. The plots show parasite genome copies (Log_10_ scale) per microlitre of DNA extracted from each culture timepoint (each culture sample was extracted into a 50 microlitre volume of DNA). The overall parasite multiplication rate (PMR) for each isolate is derived from all of the timepoint data in all of the triplicates over the six day period, applying a linear model on the Log10 density data, and this is expressed in non-logarithmic form as the multiplicative fold-change per nominal unit time of 48 hours. Other details for each isolate are shown in Supplementary Tables S1 and S2. **B**. Bar plot showing the multiplication rate (with 95% confidence intervals) of each isolate, ranked from the lowest (1.5-fold) to the highest (5.0-fold). Community-based isolates (shown with white bars) are skewed towards the left of the distribution, with generally lower multiplication rates than the hospital-based isolates (shaded bars). **C**. The distributions of parasite multiplication rates in community-based isolates (mean = 1.8-fold) and hospital-based isolates (mean = 2.9-fold) are significantly different (Mann Whitney test, P < 0.001).

Analysis of the ranked parasite multiplication rates shows that community-based isolates are skewed towards the low end of the distribution (Figure 1B). The distributions of multiplication rates in community-based isolates (mean = 1.8-fold) and hospital-based isolates (mean = 2.9-fold) are significantly different (Mann Whitney test, P < 0.001) (Figure 1C).

### Testing for correlation with peripheral blood parasitaemia and age of subjects

The estimated level of peripheral blood parasitaemia in the study subjects (Supplementary Table S1), was tested for correlation with parasite multiplication rates in the culture assays. Analysis of all samples together shows a highly significant positive correlation (Spearman’s rho = 0.49, P = 0.009) (Figure 2A). Analyses within each of the subgroups also indicate positive trends (Figure 2A). There was no significant correlation between parasite multiplication rates and age of patients, overall (Spearman’s rho = -0.09, P = 0.6) or within either subgroup (Figure 2B).

**Figure 2.**
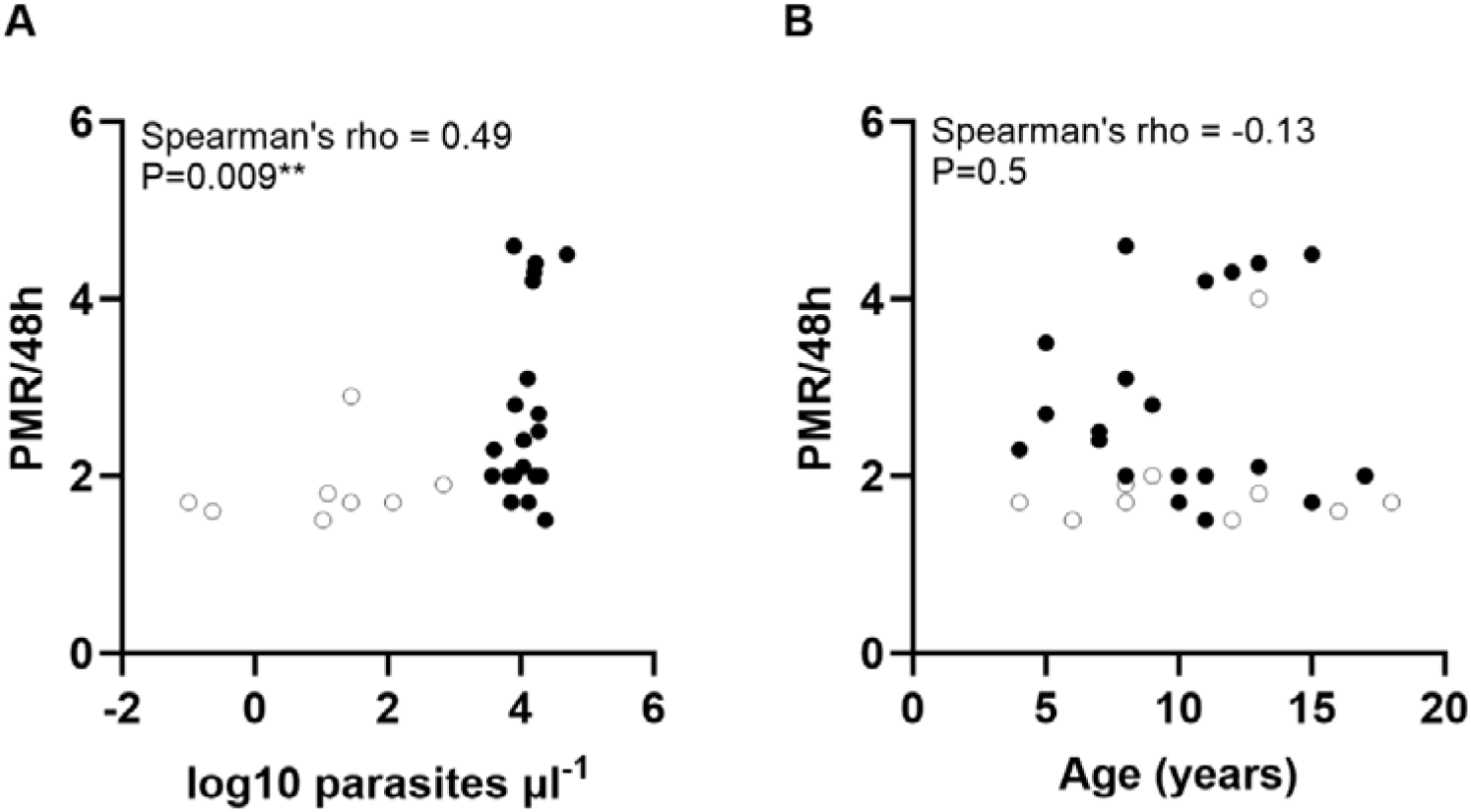
**A**. Correlation between parasite multiplication rates under exponential growth conditions in culture and the parasitaemia measured in Gambian individuals at time of original blood sampling. Open symbols show community isolates, and closed symbols show hospital isolates (measurements were available for all except six of the 34 isolates as shown in Supplementary Table S1). Analysis of all samples together shows a highly significant positive correlation (N = 28, Spearman’s rho = 0.49, P = 0.009). Although there is limited power to test correlations within each of the subgroups, these also have positive trends (community isolates N = 8, rho = 0.78, P = 0.04; hospital isolates N = 20, rho = 0.18, P = 0.44). **B**. There is no significant correlation between parasite multiplication rates and age of individuals (available for all except two of the isolates, N = 32, rho = -0.13, P = 0.5).

### Testing for correlation with genotypic mixedness and presence of gametocytes

To test which of the cultured isolates contained multiple parasite genotypes, the highly polymorphic marker loci *msp1* and *msp2* were examined (Supplementary Table S2). Overall, 74% (25 of 34 isolates) contained multiple genotypes, a proportion that was similar among the hospital-based clinical isolates (74%, 17 of 23) and the community-based isolates (73%, 8 of 11). The distribution of multiplication rates was similar for single genotype isolates (mean PMR = 2.5-fold) and multiple-genotype isolates (mean PMR = 2.5-fold) (Figure 3B). The multiple-genotype isolates each contained between two and five different genotypes (Supplementary Table S3), but there was no significant correlation between the numbers of detected genotypes and the multiplication rate of each isolate (Figure 3B).

**Figure 3.**
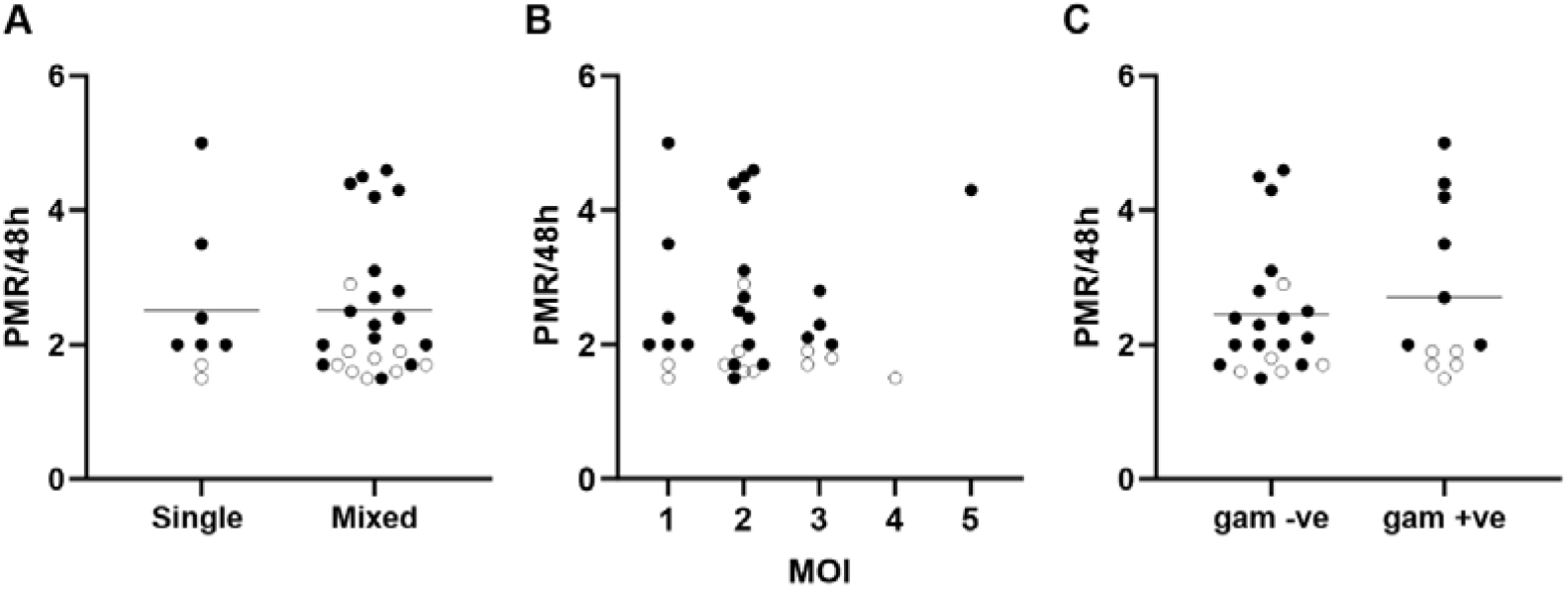
Multiplication rates of Gambian *P. falciparum* isolates under exponential growth conditions are not associated with the numbers of genotypes detected in each isolate or with the detected presence of gametocytes in culture at the time of assay. **A**. There is no significant difference in multiplication rates between single-genotype isolates (N = 8) and multiple-genotype isolates (N = 26)(mean PMR of 2.5-fold for each category, Mann-Whitney test, P = 0.9). **B**. There is no significant correlation between multiplication rates and numbers of genotypes detected within each isolate (multiplicity of infection, MOI) (Spearman’s rho = -.06, P = 0.7). Isolates were genotyped by PCR of two highly polymorphic loci (details of alleles detected in each isolate are given in Supplementary Table S3). **C**. Data on parasite stages detected in slide examinations at the time of multiplication rate assay initiation were available for 33 isolates. Isolates with any gametocytes detected had a similar multiplication rate distribution (mean = 2.7-fold, N = 12) compared to isolates without any detectable gametocytes (mean = 2.5-fold, N = 21) (Mann-Whitney test P = 0.9).

Examination of Giemsa-stained slides of parasites at the point of initiating the multiplication rate assays showed that isolates with detectable gametocytes had a similar distribution of multiplication rates (mean PMR = 2.7-fold) compared to those with no gametocytes detected (mean PMR = 2.5-fold) (Figure 3C).

## Discussion

These results indicate that intrinsic variation in *P. falciparum* multiplication potential is likely to be clinically relevant, as had been suggested by studies reporting high *ex vivo* multiplication rates in severe malaria isolates [9, 10]. Here, a significant effect of intrinsic parasite variation is suggested by comparison of isolates from clinical malaria cases attending hospital and infections sampled from the local community in The Gambia. The comparisons involved establishing continuous cultures from cryopreserved samples and performing assays under the same exponential growth conditions, showing that parasites sampled from community infections had significantly lower multiplication rates, as well as significantly lower *in vivo* parasitaemia. While not yet demonstrating causality, this supports a hypothesis that intrinsic parasite multiplication rate variation may affect whether an infection attains a high parasitaemia that increases probability of hospital presentation.

Previously, among isolates from clinical cases from a more highly endemic population in Ghana, a significantly positive correlation between multiplication rates in culture and peripheral blood parasitaemia levels at time of sampling was reported [13]. It is interesting that parasite multiplication rates in clinical isolates from The Gambia here were lower than seen in the isolates from Ghana, and the initial parasitaemia levels were also lower in Gambian patients. Although a causal relationship remains to be proven, this is consistent with a general correlation between parasite multiplication rates and *in vivo* parasitaemia as indicated within each study. Even if there were strong causality, this correlation is not expected to be absolute, as there are many other determinants of parasitaemia in the peripheral blood at a given time, and some community infections at low parasitaemia could potentially go on to become high parasitaemia infection leading to hospitalisation. The direction of causality also remains to be established, as variation in the *in vivo* blood-stage environment during infection, including differences in host erythrocytes, iron availability and inflammatory immune responses [20-22], may stimulate parasites to adjust their intrinsic growth rate phenotypes which then persist in culture.

The potential significance of mixed genotype infections has also required consideration. In a laboratory experimental model, co-infections of two strains of a rodent malaria parasite species in mice tended to be more severe than single-strain infections, although the difference was not clearly attributable to altered multiplication rates [23]. Other studies with the same model have suggested that parasites may increase their virulence during two-strain co-infections [24]. The present study showed no difference in the multiplication rates of *P. falciparum* in cultured isolates with multiple genotypes compared to those with only single genotypes detected. Analysis of clinical isolates from Ghana previously showed a trend towards higher multiplication rates in isolates containing single genotypes compared with those having multiple genotypes [13]. Therefore, these studies yield no evidence to support a hypothesis that human malaria parasites switch to higher multiplication rates in the presence of competitor genotypes.

It remains to be determined how *P. falciparum* evolves or adaptively varies its asexual multiplication rate in the blood. Although multiplication rates seen here in the clinical isolates from The Gambia were lower than those from an area of higher endemicity in Ghana, it is unknown whether such rates vary depending on local transmission conditions as previously suggested [25]. One study of parasite gene expression has suggested that there may be increased commitment to parasite sexual stages required to infect mosquitoes when overall transmission rates are lower [26]. However, the substantial variation in asexual multiplication rates in the present and previous studies is not explainable by varying proportions of parasites undergoing sexual commitment, as these are normally only a small minority of parasites per cycle [13, 27]. Consistent with this, the multiplication rate variation in the present study was not significantly associated with presence or absence of gametocytes detected in culture.

Importantly, it is not yet known which phases of the parasite asexual blood-stage cycle contribute most to the intrinsic multiplication rate variation. It has been previously noted that the moderate variation in numbers of merozoites in mature schizonts may not have a major effect [13], but merozoite invasion efficiency involving use of alternative receptors, and cell cycle duration could be significant contributors [4, 14, 28, 29]. All of the determinants may also alter when parasites are not under density-independent exponential growth conditions [4, 30]. An epidemiological perspective on the important parasite trait of variable multiplication potential highlights the need for more experimental studies to understand the genetic and cellular processes responsible.

## Methods

### Parasite sampling from malaria patients and community members

Blood samples were collected from clinical malaria cases attending Basse District Hospital in the Upper River Region of The Gambia, an area that has higher malaria incidence than the rest of the country but lower than previously experienced [17]. Patients were eligible for recruitment into the study if they were aged up to 18 years and had uncomplicated clinical malaria, testing positive for *P. falciparum* malaria by microscopical examination of a Giemsa-stained thick blood smear, and having had no antimalarial treatment within the previous month. Blood samples were also collected from community samples in villages close to Basse town, as part of other studies on the epidemiology of infection in the area [31, 32]. Written informed consent was obtained from parents or other legal guardians of all participating children, and additional assent was received from the children themselves if they were 10 years or older. Antimalarial treatment was provided according to government guidelines. Prior to treatment, venous blood samples (up to 5 ml) were collected into heparinised vacutainer tubes (BD Biosciences, CA, USA), and a proportion of each sample was cryopreserved for later analysis of parasites in culture, adding five times the volume of Glycerolyte 57 reagent (Fenwal Blood Technologies, USA) to a given packed erythrocyte volume after centrifugation. Peripheral blood parasitaemia was estimated by examination of a Giemsa-stained thick blood smear (counting the number of *P. falciparum* parasites per 200 leukocytes and multiplying by the total leukocyte count obtained by automated haematology analysis), or by using a highly sensitive quantitative PCR method [33] to assay DNA from some of the community samples that had parasitaemia levels too low for accurate estimation by slide examination (<100 μl^-1^). Approval for the original study recruitments and the parasite culture phenotyping and genotyping analyses was granted by the Joint Ethics Committee of the MRC Gambia Unit and The Gambia Government, and by the Ethics Committee of the London School of Hygiene and Tropical Medicine (reference numbers SCC 1299, SCC 1318, SCC 1476, L2014.40 and L2015.50).

### Parasite culture

Cryopreserved blood samples from sampled infections were transferred by shipment on dry ice to LSHTM where parasite culture on all samples were performed under controlled conditions within a single laboratory suite. A total of 97 blood samples (35 from hospital cases and 62 from community infections) were thawed from glycerolyte cryopreservation and *P. falciparum* parasites were cultured at 37°C using standard methods [34]. No isolates were pre-cultured before the thawing of cryopreserved blood in the laboratory. Briefly, 12% NaCl (0.5 times the sample volume) was added dropwise to each sample while gently agitating the tube to allow mixture, following which the tube was left to stand for 5 mins, then 10 times the original volume of 1.6% NaCl was added dropwise, gently agitating to allow mixture. After centrifugation for 5 min at 500 g, the supernatant was removed and cells were resuspended in the same volume of RPMI 1640 medium (Sigma-Aldrich, UK) containing 0.5% Albumax™ II (Thermo Fisher Scientific, UK). Cells were centrifuged again, supernatant removed and the pellet (comprising at least 250 μl for each sample) was resuspended at 3% haematocrit in RPMI 1640 medium supplemented with 0.5% Albumax II, under an atmosphere of 5% O_2_, 5% CO_2_, and 90% N_2_, with orbital shaking of flasks at 50 revolutions per minute. Replacement of the patients’ erythrocytes in the cultures was achieved by dilution with erythrocytes from anonymous donors every few days, erythrocytes being obtained commercially each week (Cambridge Bioscience, UK), so that after 20 days of continuous culture parasites were growing exclusively in erythrocytes from these donors, avoiding confounding from heterogeneous patient erythrocytes. Erythrocytes were screened by the commercial provider so that none were from donors with sickle cell trait. Clinical isolates were cultured in parallel in batches of up to 12 in separate flasks at the same time, so that the donor erythrocyte sources were the same for all isolates within a batch (Supplementary Table S2). After a few weeks of culture, parasite clinical isolates normally do not contain detectable mutants, which can arise and become common in some isolates after longer periods [8, 13, 35]. After at least 20 days in culture, a total of 55 isolates (31 from hospital cases and 24 from community infections) had sufficient numbers of parasites to enable initiation of exponential multiplication rate assays as described below.

### Parasite multiplication rate assays

Exponential multiplication rate assays were performed using a method previously described [12], summarised briefly as follows. Prior to each of the assays which were initiated after more than 20 days of continuous culture of the isolates, fresh blood was procured from three anonymous blood donors in the UK who had not recently taken any antimalarial drugs or travelled to a malaria endemic area, and who did not have sickle-cell trait or other known haemoglobin variants, and erythrocytes were stored at 4°C for no more than 2 days then washed immediately before use. Each of the assays for each isolate was performed in triplicate, with erythrocytes from three different donors in separate flasks (erythrocytes commercially provided by Cambridge Bioscience, UK). Asynchronous parasite cultures were diluted to 0.02% parasitaemia within each flask at the start of each assay which was conducted over six days. Every 48 hours (day 0, 2, 4, and 6) a 300μl sample of suspended culture was taken (100μl pelleted for Giemsa smear, 200μl for DNA extraction and qPCR), and culture media were replaced at these times. For each parasite isolate, a measure of exponential parasite multiplication rate was derived from all of the datapoints in all of the triplicate cultures over the six-day period, using the following procedure.

Following extraction of DNA, qPCR to measure numbers of parasite genome copies was performed using a previously described protocol targeting a highly conserved single locus (the Pfs25 gene) in the *P. falciparum* genome [12]. Analysis of parasite genome copy numbers per microlitre of extracted DNA at days 0, 2, 4 and 6 of each assay was performed, and quality control excluded any assays with less than 10 genome copy numbers per microlitre. Quality control also removed any single points where a measurement was either lower or more than 20-fold higher than that from the same well two days earlier in the assay, and any outlying points among the biological triplicates that had a greater than two-fold difference from the other replicates on the same day. Assays were retained in the final analysis if there were duplicate or triplicate biological replicate measurements remaining after the quality control steps, and if they showed a coefficient of determination of *r*^2^ > 0.75 for the multiplication rate estimates using all data points. The assay results for 34 isolates (23 from hospital cases and 11 from community infections) passed the quality control criteria. For each qualified assay, an overall parasite multiplication rate (defined as per 48-hour typical replicative cycle time) was calculated with 95% confidence intervals using a standard linear model with GraphPad PRISM. The *P. falciparum* clone 3D7 was tested in parallel as a control in all assays, consistently showing a multiplication rate of approximately 8.0 fold per 48 hours as described previously [12].

### Parasite genotyping

To test for the presence of single or multiple *P. falciparum* genotypes within each of the cultured isolates, two highly polymorphic marker loci were analysed by PCR of genomic DNA extracted from cultures. Alleles of the highly polymorphic repeat sequence regions of the *msp1* and *msp2* loci were discriminated using a nested PCR protocol with visual scoring of different allelic sizes on 1% agarose gels [36]. For each isolate, the different alleles at each locus were enumerated, and the number of alleles at the locus that had the most detected was considered as the multiplicity (minimum number of haploid parasite genotypes detected) for analysis of genotypic mixedness.

## Acknowledgements

We are grateful for the support of many colleagues at the MRC Unit in The Gambia at LSHTM where sampling was conducted in the context of ongoing studies of malaria, including Prof Umberto d’Alessandro, Mrs Haddy Nyang and Mr Simon Correa. We are grateful to patients and community members for their voluntary participation, hospital staff and community field workers, and those managing laboratory facilities and equipment as well as storage and shipment of materials. All authors contributed significantly to performing the study. Study design was undertaken by LBS, AA-N, AC and DJC; sample selection was done by LBS, AA-N, AC and DJC; laboratory experiments and assays were performed by LBS, ELE and JP; data analysis was performed by LBS and DJC; Drafting and co-ordination of the manuscript writing was performed by LBS and DJC; review and editing of the manuscript involved input from LBS, ELE, JP, AA-N, AC and DJC.

