## Supplementary Table for "Higher multiplication rates of *Plasmodium falciparum* in isolates from hospital cases compared with community infections"

**Supplementary Table S1.** Multiplication rates of *P. falciparum* in isolates from malaria cases presenting to hospital (N=23) and community-based infection samples (N=11) under exponential growth conditions in culture.

| Lab ID | Local ID | Age (yrs) | Parasitaemia in individual $\mu l^{-1}$ | Days in Culture at start of 6-day PMR assay | Parasite Multiplication Rate per 48h hours (PMR) | 95% CI of PMR estimate | $r^2$ |
|---|---|---|---|---|---|---|---|
| | | Hospital Isolates | | | | | |
| INV487 | 1299-0037.1 | 7 | 11050 | 20 | 2.4 | (2.0 - 2.8) | 0.95 |
| INV495 | 1299-0020.1 | 4 | 3929 | 20 | 2.3 | (2.1 - 2.6) | 0.98 |
| INV497 | 1299-0045.1 | 11 | 23520 | 20 | 1.5 | (1.4 - 1.7) | 0.91 |
| INV502 | 1299-0019.1 | 15 | 7165 | 20 | 1.7 | (1.2 - 2.4) | 0.84 |
| INV507 | 1299-0013.2 | 10 | 13090 | 20 | 1.7 | (1.4 - 2.1) | 0.84 |
| INV509 | 1299-0033.2 | 8 | 7838 | 20 | 2.0 | (1.6 - 2.5) | 0.91 |
| INV510 | 1299-0028.2 | 11 | 16807 | 21 | 2.0 | (1.6 - 2.5) | 0.88 |
| INV513 | 1299-0036.1 | 9 | 8290 | 21 | 2.8 | (2.2 - 3.4) | 0.94 |
| INV514 | 1299-0031.1 | 10 | 19833 | 21 | 2.0 | (1.6 - 2.7) | 0.90 |
| INV515 | 1299-0025.1 | 5 | 18715 | 21 | 2.7 | (2.4 - 3.1) | 0.98 |
| INV516 | 1299-0032.2 | 15 | 49517 | 21 | 4.5 | (3.7 - 5.4) | 0.99 |
| INV517 | 1299-0035.1 | 17 | 6716 | 21 | 2.0 | (1.8 - 2.2) | 0.99 |
| INV518 | 1299-0039.3 | 17 | 3709 | 21 | 2.0 | (1.6 - 2.5) | 0.91 |
| INV519 | 1299-0029.2 | 13 | 16679 | 21 | 4.4 | (3.8 - 5.2) | 0.99 |
| INV521 | 1299-0001.1 | - | - | 20 | 2.4 | (2.0 - 2.9) | 0.94 |
| INV522 | 1299-0056.1 | 5 | - | 20 | 3.5 | (2.9 - 4.3) | 0.96 |
| INV523 | 1299-0042.1 | - | - | 20 | 5.0 | (4.4 - 5.7) | 0.99 |
| INV524 | 1299-0014.2 | 13 | 10938 | 20 | 2.1 | (1.8 - 2.4) | 0.94 |
| INV525 | 1299-0046.1 | 8 | 12594 | 20 | 3.1 | (2.6 - 3.7) | 0.95 |
| INV528 | 1299-0055.1 | 11 | 15200 | 20 | 4.2 | (3.8 - 4.7) | 0.99 |
| INV529 | 1299-0052.2 | 8 | 7854 | 20 | 4.6 | (3.5 - 5.9) | 0.95 |
| INV530 | 1299-0050.2 | 12 | 15913 | 20 | 4.3 | (3.7 - 5.0) | 0.98 |
| INV531 | 1299-0049.1 | 7 | 18486 | 20 | 2.5 | (2.0 - 3.1) | 0.92 |
| | | Community Isolates | | | | | |
| INV341 | P0060623_03.08 | 13 | 12.4 | 38 | 1.8 | (1.6-2.0) | 0.92 |
| INV342 | P0060620 | 8 | - | 38 | 1.9 | (1.6-2.4) | 0.87 |
| INV346 | P0010112 | 6 | - | 38 | 1.5 | (1.4-1.7) | 0.88 |
| INV348 | P0020215_01.08 | 12 | 10.5 | 38 | 1.5 | (1.3-1.7) | 0.84 |
| INV349 | P0060616 | 16 | - | 38 | 1.6 | (1.3-1.9) | 0.84 |
| INV351 | P0010121 | 9 | 687 | 38 | 1.9 | (1.6-2.2) | 0.93 |
| INV363 | J0272815_23.11 | 15 | 0.1 | 38 | 1.7 | (1.5-2.0) | 0.94 |
| INV365 | K0030302_12.10 | 8 | 27.9 | 38 | 1.7 | (1.4-2.1) | 0.78 |
| INV494 | P0060606 | 13 | 28.4 | 20 | 2.9 | (2.2-3.7) | 0.89 |
| INV499 | K0364314 | 4 | 0.23 | 20 | 1.6 | (1.4-1.9) | 0.89 |
| INV508 | J0091005 | 18 | 120 | 21 | 1.7 | (1.4 – 2.2) | 0.91 |

Age in years was not recorded for two subjects although all were aged 18 years or under. Parasitaemia measurements, available for all except six of the subjects), were by thick blood film examination except where densities were <100 $\mu l^{-1}$ in which cases a highly sensitive quantitative PCR method was applied as described in the Methods.

**Table S2.** Numbers of parasite genome copies per microlitre of extracted DNA in each experimental replicate (R1 – R3 ) of each clinical isolate in six-day exponential multiplication rate assays run in four separate batches.

**Batch 1**

| Isolate ID | INV341 | | | INV342 | | | INV346 | | | INV348 | | | INV349 | | | INV351 | | | INV363 | | | INV365 | | |
|---|---|---|---|---|---|---|---|---|---|---|---|---|---|---|---|---|---|---|---|---|---|---|---|---|
| | R1 | R2 | R3 | R1 | R2 | R3 | R1 | R2 | R3 | R1 | R2 | R3 | R1 | R2 | R3 | R1 | R2 | R3 | R1 | R2 | R3 | R1 | R2 | R3 |
| Day 0 | 824 | 965 | 845 | 928 | 1093 | 1171 | 221 | 268 | 294 | 361 | 524 | 470 | 317 | - | 276 | 1313 | 1605 | 1867 | 270 | - | 433 | 889 | - | 857 |
| Day 2 | 900 | 1022 | - | - | 1370 | 1045 | 559 | 356 | 395 | 732 | 545 | 473 | 472 | - | 277 | - | - | - | 524 | - | 648 | 957 | - | 995 |
| Day 4 | 2560 | 3133 | 2553 | 3620 | 2144 | 2176 | 642 | 740 | 536 | 1181 | 1199 | 958 | 753 | - | - | 4710 | 6973 | 5685 | 1072 | - | 1024 | 2725 | - | - |
| Day 6 | 4498 | 5115 | 3920 | 9600 | 6150 | 7869 | 1195 | 983 | 705 | 1635 | 1062 | 1301 | 912 | - | 1208 | 15074 | 11316 | 6886 | 1636 | - | - | 4193 | - | 2430 |

**Batch 2**

| Isolate ID | INV487 | | | INV494 | | | INV495 | | | INV497 | | | INV499 | | | INV502 | | | INV507 | | | INV508 | | |
|---|---|---|---|---|---|---|---|---|---|---|---|---|---|---|---|---|---|---|---|---|---|---|---|---|
| | R1 | R2 | R3 | R1 | R2 | R3 | R1 | R2 | R3 | R1 | R2 | R3 | R1 | R2 | R3 | R1 | R2 | R3 | R1 | R2 | R3 | R1 | R2 | R3 |
| Day 0 | 980.7 | 586.8 | 775 | 1107 | 757 | 870 | - | 1607 | 1670 | 5695 | 4643 | 5011 | 3469 | - | 2208 | - | 1336 | 2163 | 3312 | 5038 | 6111 | - | 2384 | 2358 |
| Day 2 | - | - | - | 3498 | 6465 | 2537 | - | 3435 | 4000 | 11500 | 8708 | 7487 | 4843 | - | 4392 | - | - | - | 8377 | 9158 | 16230 | - | 4206 | 7062 |
| Day 4 | 3826 | 7230 | 6717 | 15800 | 17200 | 11000 | - | 12000 | 10000 | - | 10300 | 10200 | 5938 | - | 10200 | - | 2653 | 4276 | - | 10410 | - | - | - | - |
| Day 6 | 8863 | 9386 | 10100 | 13200 | 27700 | 21800 | - | 19500 | 19300 | 18600 | 19900 | 21900 | 11900 | - | 10400 | - | 8958 | 9809 | 23340 | 33470 | 24850 | - | 10900 | 14700 |

**Batch 3**

| Isolate ID | INV509 | | | INV510 | | | INV513 | | | INV514 | | | INV515 | | | INV516 | | | INV517 | | | INV518 | | | INV519 | | |
|---|---|---|---|---|---|---|---|---|---|---|---|---|---|---|---|---|---|---|---|---|---|---|---|---|---|---|---|
| | R1 | R2 | R3 | R1 | R2 | R3 | R1 | R2 | R3 | R1 | R2 | R3 | R1 | R2 | R3 | R1 | R2 | R3 | R1 | R2 | R3 | R1 | R2 | R3 | R1 | R2 | R3 |
| Day 0 | - | 403.7 | 546 | 475.5 | 530.9 | 510 | 410 | 512.2 | 480.4 | 422.9 | 246.9 | - | 1808 | 1739 | - | 385.4 | - | - | - | 670.5 | 675.3 | - | - | - | 744.5 | - | ND |
| Day 2 | - | 590.9 | 887 | 524.7 | 681 | 651.1 | - | 962.5 | 977.8 | 436 | 461.6 | - | 4936 | 3551 | - | 1395 | 1570 | - | - | 1236 | 1379 | 2474 | 2265 | 2298 | 3064 | 2583 | - |
| Day 4 | ND | 1853 | 1759 | 1868 | 1411 | 1466 | 1879 | 3310 | 1963 | 952.4 | 1117 | - | 11100 | 9996 | - | - | 5153 | - | ND | ND | ND | 9655 | 7457 | 7349 | 10400 | 15500 | ND |
| Day 6 | - | 2463 | 5397 | - | 5185 | - | 12500 | 10800 | - | 3060 | - | - | 42100 | 32500 | - | 37200 | 29900 | - | - | 6053 | 4749 | ND | 39600 | 22800 | 53400 | 69500 | - |

**Batch 4**

| Isolate ID | INV 521 | | | INV 522 | | | INV 523 | | | INV 524 | | | INV 525 | | | INV 528 | | | INV 529 | | | INV 530 | | | INV 531 | | |
|---|---|---|---|---|---|---|---|---|---|---|---|---|---|---|---|---|---|---|---|---|---|---|---|---|---|---|---|
| | R1 | R2 | R3 | R1 | R2 | R3 | R1 | R2 | R3 | R1 | R2 | R3 | R1 | R2 | R3 | R1 | R2 | R3 | R1 | R2 | R3 | R1 | R2 | R3 | R1 | R2 | R3 |
| Day 0 | 438 | 872 | 885 | 722.8 | 500.2 | 538.1 | 636.2 | 761.4 | - | 857.7 | 965.2 | 876.3 | 400 | 729.5 | 844.7 | 1885 | 1596 | 1642 | - | 799.7 | 773.8 | 413 | 374.1 | 556.8 | 790 | 1473 | 1788 |
| Day 2 | 1680 | - | 1371 | 1593 | 1269 | 907.8 | 5002 | 3775 | 4163 | 1688 | 1162 | 1450 | 3078 | 2499 | 2441 | - | 7630 | 6231 | 7472 | 3576 | 3484 | 1769 | 1394 | 1077 | 4942 | - | 2311 |
| Day 4 | 5909 | 4402 | 2541 | 5647 | 8336 | 6636 | 28900 | - | 20100 | 4132 | 2847 | 3064 | 9471 | 7749 | 6082 | 39400 | 36000 | 35800 | 27900 | 43500 | 18300 | 8044 | 10700 | 8312 | - | 4642 | 7721 |
| Day 6 | 9081 | 8181 | 11650 | 24700 | 20300 | - | 79800 | 91300 | 95600 | 9054 | ND | - | 26200 | 12200 | 19600 | 1E+05 | 1E+05 | - | 101000 | 54800 | 68530 | 37900 | 27800 | 27300 | 16700 | 21200 | 25500 |

Experiments were performed in 4 batches, each isolate grown for at least 20 days before assay (Batch 1 – erythrocyte donors A,B,C,D; Batch 2 – donors E,F,G; Batch 3 – donors H,I,J; Batch 4 – donors K,L,M). Each isolate was tested with three donor erythrocyte sources. ND indicates no data; dash (-) indicates measurement failed QC check.

**Supplementary Table S3.** Single and mixed genotype *P. falciparum* isolates profiled by allelic typing of polymorphic repeat regions of the *msp1* and *msp2* loci. Multiplicity of infection (MOI) estimates are used to test for correlation with parasite multiplication rates in hospital and community isolates.

| Lab ID | Mixedness | No. of *msp1* alleles | *msp1* block 2 allelic PCR products (bp) | No. of *msp2* alleles | *msp2* allelic PCR products (bp) | MOI |
|---|---|---|---|---|---|---|
| | **Hospital** | | | | | |
| INV487 | SINGLE | 1 | M-261 | 1 | IC-474 | 1 |
| INV495 | MIXED | 2 | K-234,315 | 3 | FC-309,396 IC-477 | 3 |
| INV497 | MIXED | 2 | M-216,297 | 1 | IC-651 | 2 |
| INV502 | MIXED | 2 | K-261,M-207 | 2 | FC-276, IC-471 | 2 |
| INV507 | MIXED | 2 | K-207,M-225 | 1 | IC-558 | 2 |
| INV509 | SINGLE | 1 | K-216 | 1 | IC-606 | 1 |
| INV510 | SINGLE | 1 | M-234 | 1 | IC-567 | 1 |
| INV513 | MIXED | 3 | K-342,M-234,R-156 | 2 | FC-282, IC-558 | 3 |
| INV514 | MIXED | 2 | K-351,R-156 | 2 | FC-369, IC-567 | 2 |
| INV515 | MIXED | 2 | K-351,M-243 | 1 | FC-285 | 2 |
| INV516 | MIXED | 2 | K-180,M-216 | 2 | IC-474,582 | 2 |
| INV517 | MIXED | 3 | K-306,M-225,R-156 | 2 | FC-381, IC-417 | 3 |
| INV518 | SINGLE | 1 | M-279 | 1 | FC-297 | 1 |
| INV519 | MIXED | 2 | K-306,M-288 | 1 | IC-561 | 2 |
| INV521 | MIXED | 2 | K-243,M-234 | 1 | IC-828 | 2 |
| INV522 | SINGLE | 1 | K-306 | 1 | IC-534 | 1 |
| INV523 | SINGLE | 1 | K-252 | 1 | IC-528 | 1 |
| INV524 | MIXED | 3 | K-270,M-72,R-156 | 2 | IC-426,513 | 3 |
| INV525 | MIXED | 2 | K-234,M-216 | 2 | IC-537,639 | 2 |
| INV528 | MIXED | 1 | M-252 | 2 | FC-318, IC-645 | 2 |
| INV529 | MIXED | 2 | M-225,126 | 1 | IC-420 | 2 |
| INV530 | MIXED | 5 | K-243,M-72,126,225 R-156 | 2 | FC-318, IC-552 | 5 |
| INV531 | MIXED | 2 | K-252,M-234 | 1 | IC-504 | 2 |
| | **Community** | | | | | |
| INV341 | MIXED | 2 | K-297,M-234 | 3 | FC-282,411 IC-441 | 3 |
| INV342 | MIXED | 2 | K-234,M-99 | 1 | FC-315 | 2 |
| INV346 | SINGLE | 1 | R-156 | 1 | FC-285 | 1 |
| INV348 | MIXED | 4 | K-99,108,225 M-180 | 2 | FC-447, IC-513 | 4 |
| INV349 | MIXED | 1 | R-156 | 2 | FC-295, IC-492 | 2 |
| INV351 | MIXED | 3 | K-180,216 M-234 | 1 | IC-681 | 3 |
| INV363 | SINGLE | 1 | M-252 | 1 | IC-567 | 1 |
| INV365 | MIXED | 2 | K-252,M-234 | 3 | FC-342,396,IC-552 | 2 |
| INV494 | MIXED | 2 | K-279,M-243 | 1 | IC-471 | 2 |
| INV499 | MIXED | 2 | K-297,M-243 | 2 | FC-345, IC-507 | 2 |
| INV508 | MIXED | 3 | K-225,M-252,R-156 | 2 | FC-312, IC-525 | 3 |
